# Parrot politics: social decision-making in wild parrots relies on both individual recognition and intrinsic markers

**DOI:** 10.1101/2023.10.02.560599

**Authors:** J. Penndorf, D. R. Farine, J. M. Martin, L. M. Aplin

## Abstract

1. Dominance hierarchies are generally thought to form over time via memory of repeated interactions. However, dominance hierarchies are also occasionally reported in species with fission-fusion social dynamics, where individuals may encounter large numbers of individuals, leading to incomplete social information. It it has been alternatively proposed that the complex decision-making required in these circumstances may lead to increased selection for social cognition and memory, or to the evolution of mixed strategies that rely on memory for interaction with familiars and status signals for strangers.
2. Here, test these competing hypotheses by recording social associations and aggressive interactions in a highly social, large-brained parrot, the sulphur-crested cockatoo ( Cacatua galerita). We followed 411 individuals across three neighbouring roost sites, where individuals exhibit stable dominance hierarchies within roosts, alongside a high degree of fission-fusion dynamics within roosts and regular between-roost movements.
3. We found evidence that sulphur-crested cockatoos use a two-fold social strategy when initiating or reacting to an aggression. For familiar individuals, aggressions were initiated or escalated based on rank difference. When facing less familiar individuals, decisions to interact — or escalate — were based on the relative weight, with interactions directed towards, and more likely to escalate between, individuals of similar weight.
4. Our results suggest that social knowledge remains an important determinant of aggressive interactions in highly fission-fusion systems, but that individuals can dynamically incorporate other potential cues of competitive ability when knowledge is lacking.

## Introduction

Contests over limited resources are a fact of life for almost all social species (Ward and Webster, 2016), and can potentially be extremely costly for the animals involved. Various mechanisms should therefore evolve that allow animals to better assess competitors and more accurately target a subset of individuals to engage with, decreasing the overall level of aggression (Arnott and Elwood, 2009; Dehnen et al., 2022). There have been two main mechanisms proposed. The first is the so-called *badge-of-status* (reviewed by Santos et al., 2011). Here, individuals exhibit honest signals that correlate with fighting ability, allowing opponents to assess their probability of winning without any need for social recognition or memory. For example, house sparrows (*Passer domesticus*) display a variably sized bib patch (Liker and Barta, 2001; though see Sanchez-Tojar et al., 2018), great tits (*Parus major*) have a prominent black breast-stripe that varies in width (Järvi and Bakken, 1984; but see Wilson, 1992), and some wasp species display their quality in their facial patterning (Tibbetts and Lindsay, 2008). The second main mechanism to decrease cost of contents is the formation of relatively stable dominance hierarchies, which are primarily shaped by memory of repeated interactions and potentially by observations of the interactions amongst others (Cheney & Seyfarth, 2008). Hierarchies can either be based entirely on social recognition and memory (Barnard and Burk, 1979; Tibbetts and Dale, 2007), or a combination of social recognition and some intrinsic and predictable markers (e.g. age or sex; Chiarati et al., 2010).

Within stable groups, dominance hierarchies will allow the most precise assessment of the status of group co-members (Drews, 1993). Conversely, in open societies a badge-of-status provides an assessment mechanism, that, while not as accurate, does not require individual recognition (Sheehan and Bergman, 2016). However is now increasing evidence that many species maintain complex differentiated social networks in which groups exhibit fission-fusion dynamics, or interact with individuals from other groups (Papageorgiou et al., 2019; Papageorgiou & Farine, 2021; Camerlenghi et al., 2022). For example, many corvids and parrots exhibit classic *communal roosts*, where individuals sleep at stable roosts with potentially hundreds of individuals, forage in smaller fission-fusion subgroups, and occasionally also interact with members of other neighbouring roosts (Loretto et al., 2017; Aplin et al., 2021). There have been various arguments for what strategies individuals may use to mediate aggressive interactions under these circumstances. First, the ability to segregate into sub-groups with flexible membership may decrease the cost of competition to the extent that the need for stable hierarchies or a badge-of-status is weakened (de Silva et al., 2017) or absent (Boehm, 1999). Alternatively, complex social cognition may allow individuals to remember large numbers of individuals and interactions to maintain hierarchies (as argued for ravens: Boeckle and Bugnyar, 2012). A third understudied possibility is that individuals could use mixed strategies, with memory-based dominance hierarchies determining interactions among familiar individuals (e.g. at the roost-, or group-level), and with individuals either (i) deploying no assessment strategies towards relative strangers, (ii) assessing strangers by using a badge-of-status, or (iii) assessing strangers using a set of cues to their state. The potential existence of mixed strategies could help explain how animals manage the cognitive load associated with living in large-scale societies.

In Sheehan and Bergman (2016), for example, the authors suggested that, after dispersal, juveniles should rely on quality signals when interacting with unfamiliar individuals, but once juveniles are settled, interactions reoccur, which should favor social recognition over quality signals. Empirical evidence of mixed strategies was provided by two recent studies investigating the potential for context dependent use of badge-of-status in captive birds (Vedder et al., 2010; Chaine et al., 2011; Chaine et al., 2018). By experimentally manipulating colourful crown patches in blue tits (*Cyanistes caeruleus*) and golden-crowned sparrows (*Zonotrichia atricapilla*) and then inducing contests, the authors found that the crown patch size mediated contest outcomes amongst strangers, but was less important in determining contest outcomes in birds caught the same area (Vedder et al., 2010; Chaine et al., 2011; Chaine et al., 2018). These results suggest that animals may use individual recognition to mediate interactions with familiar individuals, and status symbols in interactions with relative strangers (Chaine et al., 2018). However, these studies inferred familiarity from spatial proximity, and did not explore whether individuals formed dominance hierarchies within familiar flocks. To the best of our knowledge such mixed-strategies are yet to be reported in natural interactions, or observed in the wild.

Here, we investigate patterns of aggressive interactions within and between three neighbouring roost groups of wild sulphur-crested cockatoos (*Cacatua galerita*). Sulphur-crested cockatoos (SC-cockatoos) are large, white, sexually monomorphic parrots that exhibit no clear plumage variation between individuals that might represent a badge-of-status (Figure 1). Like many parrots, SC-cockatoos form year-round communal roosts of 100-1000 birds (Aplin et al., 2021; Penndorf et al., 2022). Within these roosts, SC-cockatoos display highly fluid fission-fusion dynamics, with flock size varying between 2 and 500 individuals (Noske et al., 1982; Styche, 2000). In addition, birds also regularly forage with individuals of different roosts, and occasionally engage in between-roost movements (Aplin et al., 2021; Penndorf et al., 2022). Despite these larger dynamics, individuals maintain stable, long-term relationships beyond the pair bond that are strongly suggestive of social recognition, including with birds from other roosts (Aplin et al., 2021; Penndorf et al., 2022). Our previous work has also shown that SC-cockatoos maintain dominance hierarchies at the level of the roosting community, with these hierarchies stable over extended periods of at least 3 years, and likely to be memory based (Penndorf et al., 2024). Given this social structure, we hypothesised that individuals would use mixed strategies; using dominance hierarchies to determine interaction decisions at the roost level, but deploying either no assessment strategies or cue-based assessment when interacting with individuals beyond this social group.

**Figure 1:**
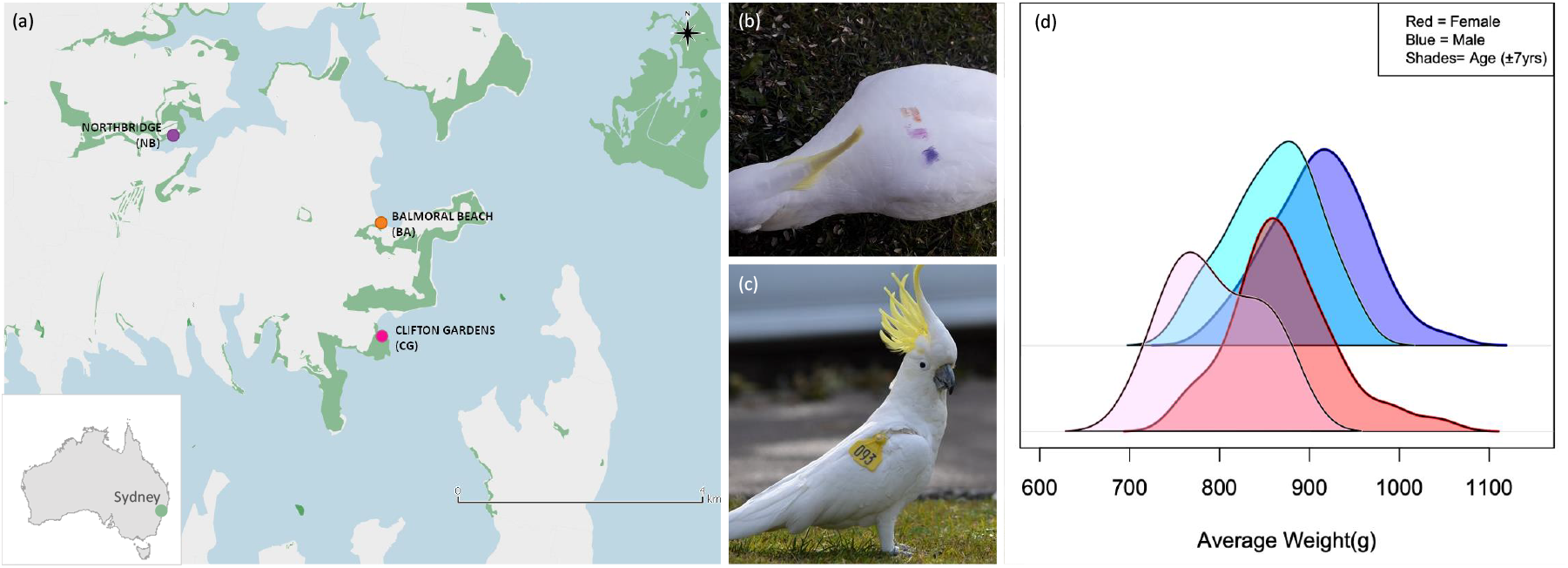
Study locations and marked birds and their respective weights. (a) Map of the three roosting communities included in the study, (b) Example of marking with temporary dye, Violet Pink Orange - horizontal, that remains visible for 3-4 months. (c) Example of wing-tagged bird — 093 Albie. (d) Distribution of average weights of individuals in grams, across age and sex classes. Dark blue are adult males (>7yrs, n=137), light blue are juvenile males (<7yrs, n=49), dark red are adult females (>7 yrs, n=116), and pink are juvenile females(<7yrs, n=30).

**Figure 2:**
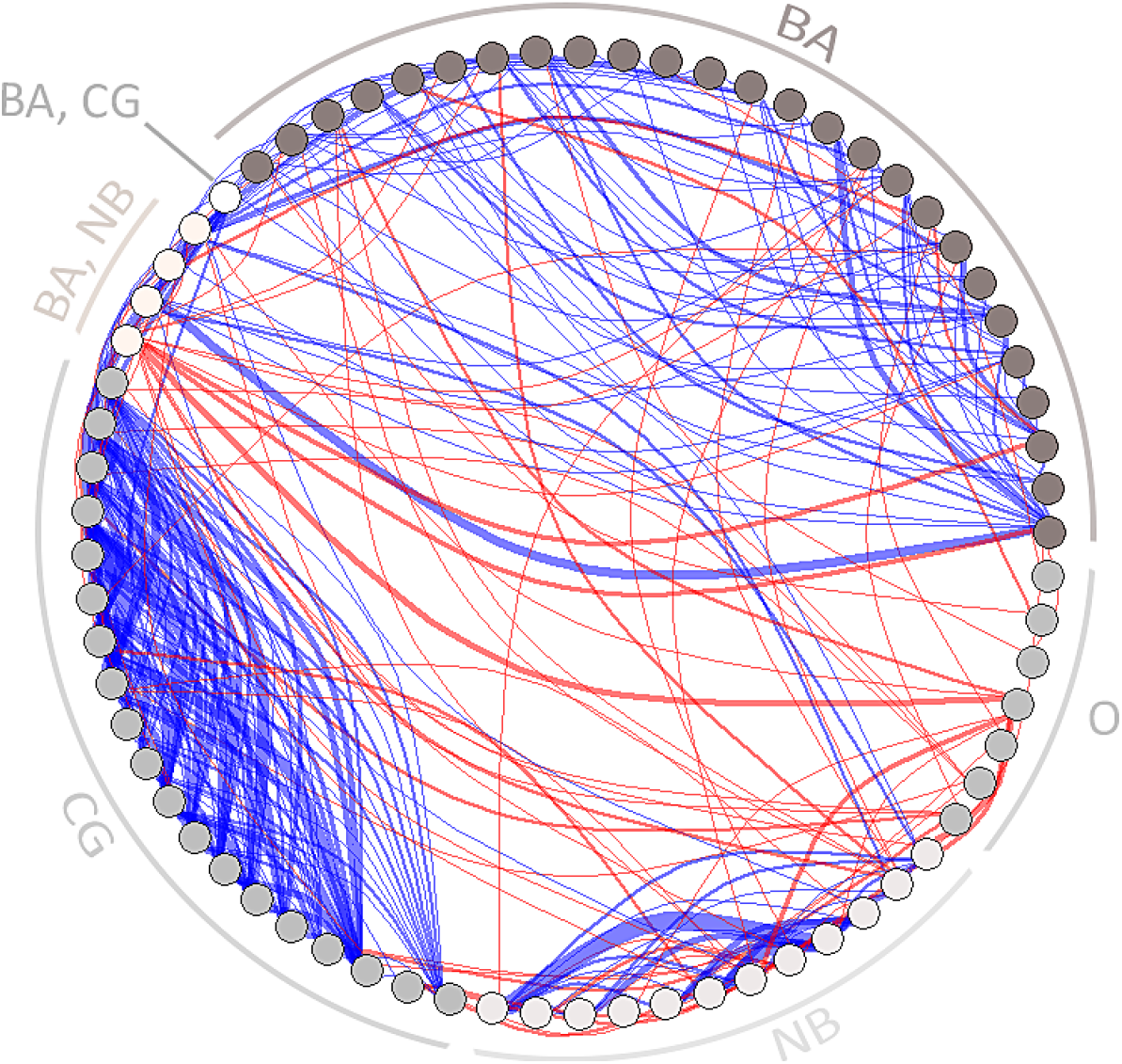
Aggressive interactions between males. Nodes are shaded by the familiarity of the individual with specific social environment(s), indicated as label for each node (BA, CG, NB). Individuals that were transient to all three sites are marked as other (O) roost-membership. Edges are coloured depending on the familiarity with the social environment. Blue edges represent interactions between familiar individuals. Red edges represent interactions between individuals from different social environments.

## Methods

### Study population

The study was conducted in an urban population of SC-cockatoos in Sydney, Australia, an area within the native range of the species. SC-cockatoos in this area use communal sleeping roosts of up to 200 individuals. These roosts are maintained year round and likely over many decades, and consist of non-breeding and breeding birds, with breeding individuals nesting in tree-hollows in close proximity to the main roost (Aplin et al., 2021). In this area, roosts tend to located 1.5-5km apart, with birds generally foraging in an approximate 3km radius around the roost (Aplin et al., 2021), and returning to the roost regularly throughout the day. Our study was focused on three similar sized neighbouring roost-sites in north Sydney, located at Balmoral Beach (BA), Clifton Gardens (CG), and Northbridge (NB): Figure 1a, Table S1.

At each of the three roost sites, birds were first habituated to the researchers and then marked using non-toxic dye (Marabu Fashion Spray, MARABU GmbH) using methods detailed in Penndorf et al. (2022). Each individual was marked with an unique combination of one to four coloured dots, applied with sponges on the middle of the back (Figure 1b). As part of a parallel project run at the same time, birds were also habituated and marked at two roost-sites outside of the focal study area (Table S1), in Manly (MA) and the Royal Botanic Gardens (BG; Table S1). In addition to the paint-marked birds, 144 birds equipped with wingtags as part of the ongoing citizen science study*Big City Birds* (Davis et al., 2017; Aplin et al., 2021, Figure 1c) regularly visited the three roosting locations.

Overall, 411 individually identifiable birds were included in the study across the three roost locations, which we estimate to be 95% (CI=0.92-0.98) of the individuals at each site (Table S1). Of these 411 birds, 373 were paint-marked, 28 were wing-tagged, and 11 had distinctive physical features that meant marking was not required (e.g., healed injuries): Table S1. Age (juveniles: <7 years, adults: >7 years) and sex of birds were assessed by eye-colour (Berry, 1981). Additionally, feathers were collected for a parallel study, and molecular sexing was used to sex juveniles (BA: n=68, CG: n=55, NB: n=41, Penndorf et al., 2022). Finally, we recorded the weight of individuals by training them to step on a flat scale that read at 1g accuracy in exchange for a food reward (e.g. sunflower seeds). This resulted in 214 birds being weighed 1 to 17 times each (mean: 4.3) over the 4 months across the three primary roost-sites (BA, CG, NB). Weight was highly repeatable within individual (0.78, 95% CI=0.72-0.82, R-package *rptR*, Stoffel et al., 2017, no. bootstraps=1000) and ranged from 717g to 1054g. Males tended to weigh more than females, and adults more than juveniles (Figure 1c).

### Social data collection

We recorded social associations and interactions over two 10-day periods from July 8th - 20th and September 19th - October 2nd 2019. Behavioural observations were collected daily for 3 hour (July) or 2.5 hour (September), during which periods birds were encouraged to forage on the ground by scattering small amounts of sunflower seeds over an approximate 385-500*m*^2^ area of grass close to the roost location (300-680m distance).

During each sampling period, presence scans (Altmann, 1974) were conducted every 10 minutes to record the identity of all birds present. We defined present as being identifiable within the park. Additionnally, we used all occurrence sampling (Altmann, 1974) to record aggressive interactions, recording the time, the identities of the individuals involved, as well as the sequence of the interaction. All aggressive behaviours recorded in the study, as well as their definitions, can be found in Table **??**. If several dyads interacted very close in time, we prioritized recording the identities of winners and losers.

### Dominance hierarchies

Given that the hierarchies of both observation periods were highly correlated (Penndorf et al., 2024), we combined the interaction data from both observation periods. We then calculated a separate dominance hierarchy for each of the three roost-sites using randomized Elo-ratings (R-package *aniDom*, Sánchez-Tójar et al., 2018; sigma=1/300, K=200, randomisations=10,000) and included only individuals with ten or more agonistic interactions at a given roost-site location (BA: N_*ind*_=126; CG: N_*ind*_=93; NB: N_*ind*_=74; Sánchez-Tójar et al., 2018).

To test whether rank was correlated with the intrinsic factors of age, sex and weight, we ran a generalized linear mixed model using the R-package *lme4* (Bates et al., 2015). We tested whether standardized rank was predicted by the average weight of an individual, as well as age and sex. Since individuals could appear in the hierarchies of several sites (26 individuals present in two hierarchies, two individuals present in three hierarchies), we also included site and individual ID as random variables.

### Social decision-making

Decisions about interactions were made at two levels. First, an individual (hereafter *initiator*) decides whether to engage in an aggressive interaction. Second, the individual that is being aggressed (hereafter *receiver*) chooses whether to retreat or reciprocate. Decision-making in both cases is based either on an assessment of the opponent’s resource holding potential (RHP, Green and Patek, 2018) or on memory, including prior experience or knowledge of relative rank (Chaine et al., 2018).

We hypothesised that in the absence of local knowledge, individuals would use assessment of RHP to decide on whether or not to engage in an aggressive interaction. We further hypothesised that when individuals had knowledge about the social environment, they would instead base their decision-making on the dominance hierarchy. In order to test these hypotheses, we considered two instances of decision making: the decision of the initiator about whom to aggress, and the decision of the receiver about how to react to the aggression.

As aggressive interactions involving females were relatively rare, we restricted interactions to male-male dyads only. As a proxy for RHP, we used an individual’s average weight, which is likely to be correlated to overall body size. As a proxy for familiarity, we considered an individuals familiar with the social environment at the site if they were present in at least 5% of the scans conducted at the site. Else, individuals present in fewer than 5% of scans at the site were considered unfamiliar with the social environment.

### Deciding whom to aggress

To test which cues SC-cockatoos are most likely to base their decisions to initiate an aggressive interaction upon, we used a two step approach. First, we tested the influence of weight on decision-making. For this, the data was divided into two groups: 1) interactions where the initiator had no social knowledge, and 2) interactions where the initiator had social knowledge. In a second step, we focused only on interactions where the initiator had social knowledge to test whether familiar individuals use an alternative, rank-based, interaction strategy.

In order to test the strategic use of aggressions for each of the above described scenarios, we adapted the method developed by Hobson et al. (2021) and modified by Dehnen et al. (2022), consisting of several steps. First, we calculated the difference between individuals in weight or rank as:

- (weight initiator) - (weight receiver) for the analysis on weight
- (rank initiator) - (rank receiver) for the analysis on rank, where dominance ranks were calculated using randomized elo-rating (R-package *aniDom*, Sánchez-Tójar et al., 2018; sigma: 1/300, K: 200, randomisations: 10,000).

The interaction data were used to count the number of observed directed interactions across each pairwise combination of individuals. To generate expected interaction patterns, we implemented a permutation procedure, where each iteration (n=1,000) randomly selected one interaction, and then randomly selected a new receiver of the interaction among all individuals present (i.e. individuals present in the scan just before and just after the interaction, including the original receiver but not the initiator). Permutations were limited to interactions where at least three were present at the site at the time of the interaction (including the original initiator and receiver).

We ran 10,000 permutations before calculating the difference between observed and permuted datasets. If the observed tendency to interact follows a random pattern, the confidence intervals should overlap zero. A positive or negative value that does not overlap zero indicates that interactions between individuals of that rank difference occurred more or less often than expected by chance. As interaction strategies towards individuals positioned higher or lower than oneself in the hierarchy may differ (Dehnen et al., 2022), we fitted separate smoothing splines for positive and negative rank differences (*smooth*.*spline* R-function, d.f.=3, lambda=0.04).

### Analysis of escalations

To examine the decision of the receiver, we identified escalation events. These were defined as interaction sequences in which the receiver responded aggressively towards the initiator of the interaction, independent of who eventually won the interaction. Across all roost sites, this resulted in 3,845 interactions between 396 individuals (Table S1).

We first tested the influence of weight and familiarity on the likelihood of an interaction to escalate. Therefore, we tested whether escalation events were predicted by the absolute weight difference (in grams), the knowledge status of both receiver and initiator, whether the initiator was of higher/lower weight than the receiver, as well as the interactions between all terms.

In a second step, we focused only on a subset of interactions, where both individuals were knowledgeable of the social environment to test the influence of rank difference on the likelihood of escalation. Therefore, we tested whether escalation events were predicted by the the absolute rank difference the knowledge status of both receiver and initiator, whether the initiator was of higher/lower rank than the receiver, as well as the interactions between all terms.

For both analyses, we constructed binomial GLMMs (package *lme4*, Bates et al., 2015) that included the base predictor for each hypothesis (absolute weight or rank differences) and all possible combinations of additional predictors (familiarity and whether the initiator was heavier for the weight hypothesis, and whether the initiator was heavier or higher in rank for the rank hypothesis) that could contribute to an escalation. We then compared models using AIC to identify which set of additional predictors were important in modulating the role of the hypothesised heuristic.

## Results

The roost-level hierarchies included between 74 and 126 individuals (i.e. birds with ≥ 10 observed interactions at a specific location; Table S1). From these 265 individuals we had known weights for 176. We recorded 6,402 aggressive interactions across all birds at each of the three roost sites, of which 5,447 were between individuals in the same dominance hierarchy (n = 265 birds). Of these, 3,859 observations had a full sequence starting with information about the initiator and receiver (see Table S1 for a subset of birds per site).

### Predictors of dominance rank

Overall, standardized weight was not significantly correlated with rank (*β*= −0.0002; P=0.570). When repeating the same analysis within males only, we found again that weight did not predict dominance rank (*β*=-0.0005, P=0.63). However, as expected, males tended to be more dominant than females. Given this, and given that most interactions involved males, the subsequent analyses were conducted only within males.

### Social decision making

#### Initiation of interactions

If the initiator was not knowledgeable of the social environment, they preferentially targeted other individuals close in weight (−170g - 200g; Figure S1a). By contrast, interactions initiated by knowledgeable individuals (present in >5% of the scans at that specific location), showed no significant influence of weight on the decision to initiate an aggressive interaction, though there was a tendency of individuals to initiate interactions with others that were slightly lighter (<80g, Figure S1b). Rather, when individuals were knowledgeable of the social environment, relative rank difference was a better predictor of social interactions. In this case, individuals directed aggressive interactions towards lower ranking individuals, while avoiding aggressing those higher in rank (Figure S1c).

#### Escalation of interactions

The model of escalation most supported by the data included an interaction between absolute weight difference and familiarity, plus a term capturing whether the initiator was higher or lower in weight. This suggests that familiar and unfamiliar individuals had different tendencies to escalate interactions for a given difference in weight. Specifically, we found that escalation events in our dataset were best predicted by an interaction between weight difference and familiarity, and whether the initiator was heavier than the receiver of the interaction (Figure 3a). That is, if one individual in the dyad had no social knowledge, then interactions were much more likely to escalate if individuals were close in weight. This probability of escalation was also higher if the receiver was heavier than the initiator (Figure 3a). If both individuals were familiar with the social environment, escalations were also more likely if the receiver was heavier, independent of the weight difference. There was a possible trend towards a higher probability of escalation when the weight difference was large (Figure 3a), however this was entirely driven by two interactions between the same dyad.

**Figure 3:**
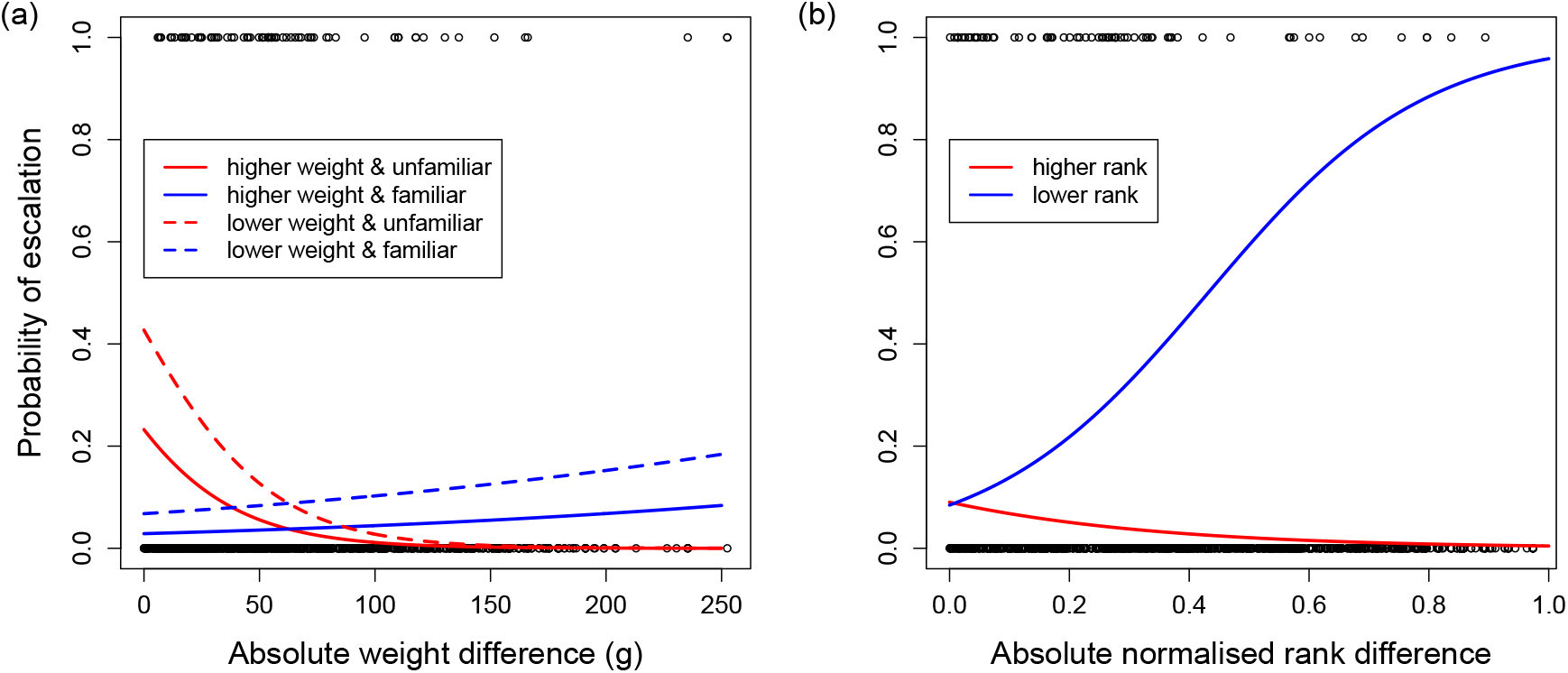
Probability for interactions to escalate depending on differences in weight and rank, and whether birds were familiar or not. (a) We found that escalations were generally more likely if the initiator had a lower weight than the receiver (dashed lines versus solid lines). In the absence of social knowledge (red lines; unfamiliar), we further found that escalations were more likely if the two individuals were closer in weight. (b) Escalation in interactions between knowledgeable individuals were only likely to occur if the initiator was lower in rank (blue line), with this probability increasing as this difference in rank increased. Line colours are labelled from the perspective of the initiator.

Second, we focused on a subset of the data containing only interactions between individuals familiar with the social environment to test whether rank differences predict escalations of aggressive interactions. For the effect of rank difference, the model most supported by the data included an interaction between absolute rank difference and whether the initiator was ranked higher than the receiver. Specifically, we found that if the initiator was ranked higher, interactions were very unlikely to escalate, and escalations only occurred if individuals were close in rank (Figure 3b). If, on the other hand, the initiator was ranked below the receiver in the hierarchy, the probability of escalation was high and increased with increasing rank difference (Figure 3b).

## Discussion

We found that SC-cockatoos used mixed decision-making strategies in their aggressive interactions. First, for dyadic interactions where at least one individual was not knowledgeable of the social environment (e.g. an individual joining a foraging flock at a different roost site), individuals disproportionately directed aggression towards those close above and below themselves in weight, peaking at 90g heavier. For a male of average weight (900g), this difference represents 10% of body weight, a difference that may not be ascertainable visually, suggesting that individuals are challenging opponents of similar overall body size. Within these interactions, the probability of escalation also depended on whether the individuals were close in weight, with this probability being higher if the receiver was higher in weight. This contrasted the interactions among familiar individuals. Here, weight had little correspondence with whom individuals directed aggression towards, and only affected the probability of escalations when the initiator was considerably lower in weight than their aggressor.

Second, within regularly interacting individuals, we found that individuals were less likely to direct aggression towards those above them in the hierarchy than expected by chance, and were more likely to direct aggression towards those below them, with most aggressions occurring at at 15-45% rank difference. Escalation to received aggression was then only likely to occur when the receiver was much higher ranked than their aggressor. Together with our results on interactions between less familiar individuals, these results provide evidence that SC-cockatoos are engaging in a mixed strategy that depends on the familiarity between interacting individuals. When facing less familiar individuals, decisions to interact — or escalate — were based on the relative weight difference. Conversely, social decisions between familiar individuals were based on social recognition and memory, with individuals generally aware of relative rank differences between themselves and familiar others.

### Interactions with strangers: using proxies for resource holding potential

When interacting with non-residents, SC-cockatoos used body size (weight) in their aggressive decision-making. This suggests that: (i) there are limits to social cognition in SC-cockatoos, with individuals unable to retain (or not using) memory of relative rank for individuals outside of their group, despite being members of nearby roosts and therefore potentially having interacted previously to our observation period, and (ii) SC-cockatoos are assessing these individuals using a set of cues to their state, including sex and body size. This finding contributes to the growing evidence of mixed strategies in systems with fission-fusion dynamics (Vedder et al., 2010; Chaine et al., 2011; Chaine et al., 2018), but extends this body of work in two important ways. First, it demonstrates the existence of such mixed strategies in wild, naturally interacting individuals, and where stable, quasilinear dominance hierarchies are maintained within familiar individuals. Second, it gives evidence for the existence of such strategies even in the absence of clear status signals. This is particularly intriguing, as it has been previously suggested that the use of social recognition could constrain the evolution of open social systems, while the use of quality signals may facilitate the evolution of open social systems (Sheehan & Bergman, 2016). It further suggests that social cognition cannot be assumed in systems without such signals of quality.

While we show an effect of assessment of body size (as estimated by weight) on decision-making, we cannot exclude the possibility that weight is correlated to a yet undescribed status signal in this species. One possibility could be crest length or colour, given its importance for signalling across Cacatuidae (Liévin-Bazin et al., 2018; Bertin et al., 2020). Another potential status signal might be UV-reflective plumage (Pearn et al., 2001; Berg and Bennett, 2010). Such fluorescent patches are found in a majority of parrot species (140 out of 143 species examined by Mullen and Pohland, 2008). SC-cockatoos have yellow feathers on their cheek, under-wing, and crests that exhibit ultraviolet fluorescence (Figure 1b) (Mullen and Pohland, 2008). The function of this fluorescence in SC-cockatoos has not been studied. However, UV-reflective plumage is used to inform mate choice in Budgerigars (Pearn et al., 2001), and it is largely thought to be a sexual signal (Berg and Bennett, 2010; Delhey et al., 2017). Whether it could also signal status and/or condition in parrots warrants further investigation. That said, based on our recorded interaction sequences it seems unlikely to be the primary cue or signal. We observed no obvious movement or display of cheek patches during aggressive interactions. Crest movements are often induced by surprising or alarming stimuli (Liévin-Bazin et al., 2018), and are utilised in breeding displays at nest hollows (*personal observation*). These observations are further strengthened by our data, as only 16% of all aggressive interactions included crest-erection (15% of interactions involving familiar individuals, and 19% of interactions involving unfamiliar individuals).

### Interactions with familiar individuals: relying on social cognition instead of proxies for rank

Our results from interactions between relative strangers suggest that SC-cockatoos are able to assess body size (for which we used weight as a proxy), and perhaps use this as a signal of RHP. Yet within roosts, weight did not determine dominance rank. This may seem surprising, given that: (i) that flocks of wild SC-cockatoos have been previously suggested to be too large to form effective dominance hierarchies (Noske et al., 1982 — but see Penndorf et al., 2024), (ii) the high degree of fission-fusion dynamics in this species means individuals may encounter hundreds of others over days and weeks(Aplin et al., 2021; Penndorf et al., 2022), and iii) that weight does partly influence decisions to initiate and/or escalate aggressive interactions. One potential explanation is that weight-based hierarchies tend to be restricted in terms of maximum group size when weight differences become too small to assess (Ang and Manica, 2010). Groups of SC-cockatoos may therefore be too large for primarily weight-based hierarchies to be useful. By contrast, SC-cockatoos must have the cognitive ability to learn and remember their position in the dominance hierarchy; noting that this hierarchy is highly stable over time (Penndorf et al., 2024). Previous work has suggested that SC-cockatoos exhibit preferred social associates (Penndorf et al., 2022), and form stable long-term relationships (Aplin et al., 2021). This supports other work in other parrot species that is also suggestive of extensive individual recognition (Wanker et al., 1998; Buhrman-Deever et al., 2008). Overall, this suggests that SC-cockatoos may possess sufficient memory of past interactions to allow them to track dominance relationships in large social groups (Hobson, 2020). Given that memory-based dominance hierarchies are predicted to be the most reliable way to assess status, this would lead individuals to rely on social cognition within roosts, only using a cues to body condition when this is not possible due to complete unfamiliarity or cognitive constraints (in the case of rare interaction partners).

## Conclusions

Our previous work has shown that wild SC-cockatoos form stable long-term relationships (Aplin et al., 2021), and maintain some social relationships even after dispersal into different roost groups (Penndorf et al., 2022). This suggest that SC-cockatoos possess extensive memory of past interactions — an ability displayed by other large-brained bird species that exhibit fission-fusion dynamics, such as common ravens (*Corvus corax* Boeckle and Bugnyar, 2012). This social cognition is further exhibited within roost groups, with birds maintaining impressively large dominance hierarchies that are stable over time (Penndorf et al., 2024). However when this knowledge is lacking (i.e. when facing less familiar individuals) aggressive decisions in SC-cockatoos to interact — or escalate — are based on the relative weight difference. This implies that, even with good memory and no signals of quality to use as proxy for RHP, a mixed strategy that depends on the frequency of aggressive encounters might be more adaptive in open societies than a sole reliance on social recognition and memory, perhaps because it may reduce errors from inaccurate recollection or outdated information. Taken together, our results suggest that social knowledge remains an important determinant of aggressive interactions, even under such high fission-fusion dynamics, but that individuals can flexibly incorporate other potential cues of competitive ability when recent knowledge is lacking.

## Acknowledgements

We acknowledge the Gamaragal and Gadigal people of the Eora Nation as the Traditional Custodians of the Land on which this study was conducted. All procedures were approved by the ACEC (ACEC Project No. 19/2107), and were conducted under a NSW Scientific License (SL100107). This work was supported by the Swiss State Secretariat for Education, Research and Innovation (SERI) under contract number MB22.00056. Part of the work was also supported by a National Geographic Grant NGS-59762R-19 to LMA, and by funding from the Max Planck Society to JP and LMA.

## Supplementary material

**Table S1:**
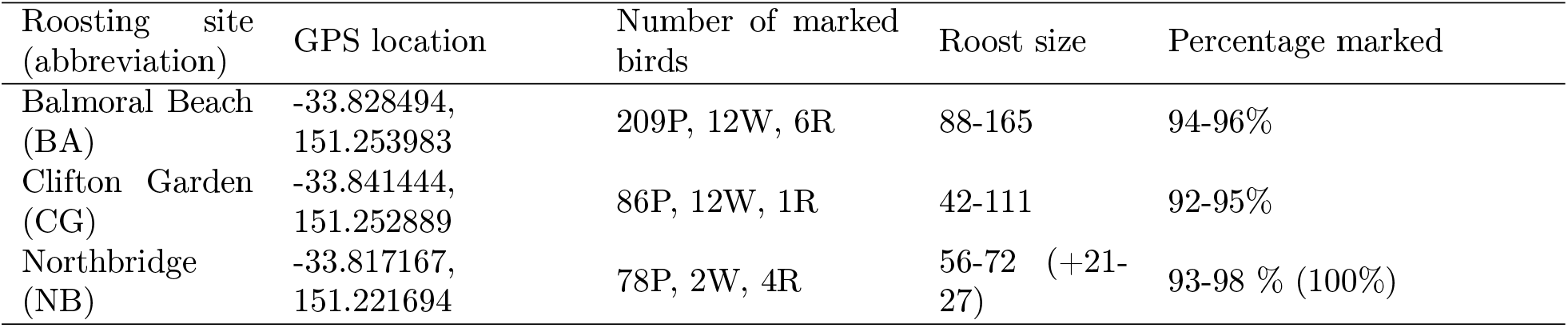
Location of the three study roosts and the estimated number of SC-Cockatoos at each site. The two letter code between parentheses represents the abbreviation for each roost site. At each site, we conducted roost counts by attracting birds to the ground after wakening, and recording the number and identities of all birds present. These roost counts were then used to estimated the number of birds at each site, as well as the percentage of marked birds. As individuals regularly visit other different sites, the number of birds marked at one specific site can be larger than the number of individuals roosting at this location. P: paintmarked; W: wingtagged; R: recognizable feature. For Northbridge, the number between brackets represent the number and percentage of marked individuals of the satellite roost.

**Table S2:**
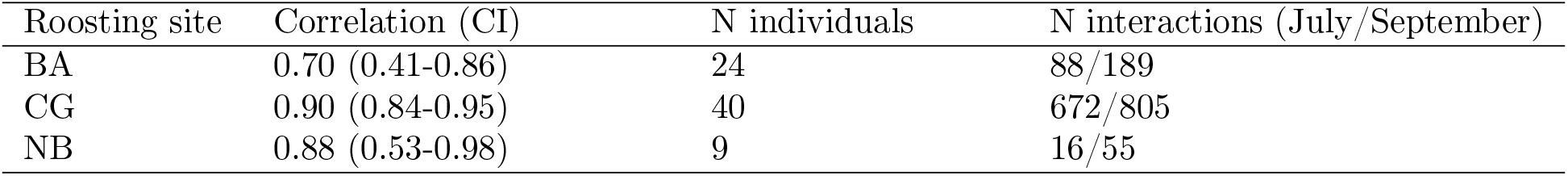
Correlations between SC-cockatoo hierarchies recorded at each roosting site during the two observation periods (July & September 2019)

**Table S3:**
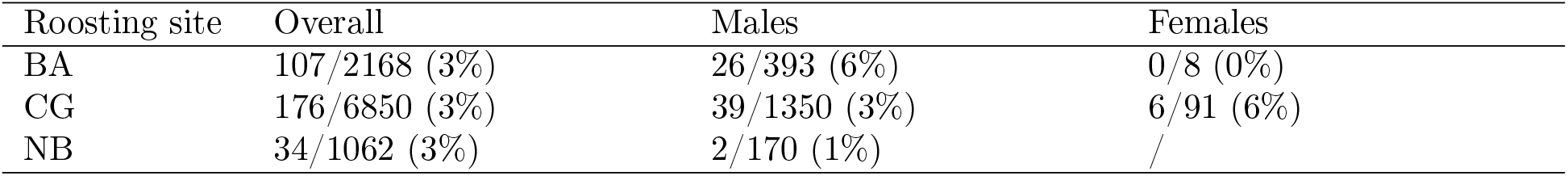
Number of SC-cockatoo cyclic triads compared to the total number of complete triads. The number of each type of triad was determined on an outcome matrix, using igraph (Csardi and Nepusz, 2006).

**Figure S1:**
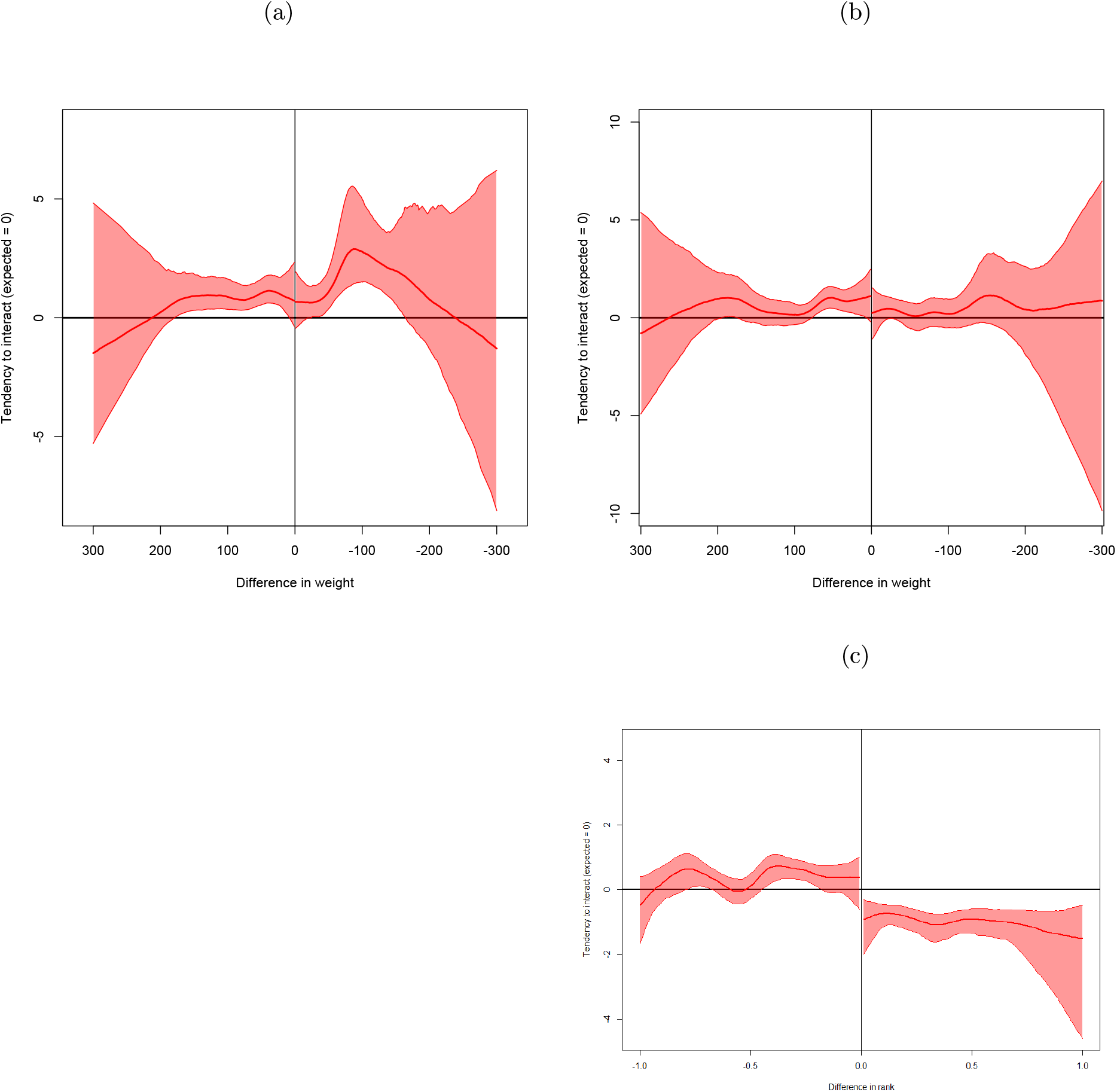
Tendency to initiate aggressive interactions, plotted by weight difference (top row - a, b) or rank difference (bottom row - c), depending on whether the initiator was considered not knowledgeable (left column) or knowledgeable (right column) of the social environment at the site of interaction. The thick line on each graph represents the median tendency to interact. The shaded areas represent the 95% confidence interval of the estimated interaction tendencies. If the tendency to interact follows a random pattern, the confidence intervals overlap with zero. A positive or negative value indicates that interactions between individuals of this specific rank difference occurred more or less often than expected by chance.

