## Supplementary material for "Parrot politics: social decision-making in wild parrots relies on both individual recognition and intrinsic markers"

### 1 Methods

#### Transitivity of the hierarchy

In order to assess the transitivity of the hierarchy, we used a directed 1/0 matrix, based on the majority of wins (McDonald and Shizuka, 2013), with the modification that when two individuals won the same number of contests against each other, they both received scores of 1 (Shizuka and McDonald, 2015). This produced a directed network where a dominant-subordinate relation was represented by an edge directed from the dominant to the subordinate individual. We used the package *compete* (Curley, 2016) to infer transitivity of dominance relationships both at group level, and within each sex. Emphasis was on the proportion of transitive triangles relative to all triangles (Pt, defined by McDonald and Shizuka, 2013) is given by:

$$P_t = \frac{N_{transitive}}{N_{transitive} + N_{cycle}}$$

where $N_{transitive}$ is the number of transitive dyads, and $N_{cycle}$ is the number of cyclic triangles. In a random network, $P_t$ is expected to be 0.75 (McDonald and Shizuka, 2013). Therefore, transitivity was scaled so that it runs from 0 for the random expectation to 1 (all triangles are transitive), by the following triangle transitivity metric ($t_{tri}$):

$$t_{tri} = 4 \times (P_t - 0.75)$$

Table S1: Location of the three study roosts and the estimated number of SC-Cockatoos at each site. The two letter code between parentheses represents the abbreviation for each roost site. At each site, we conducted roost counts by attracting birds to the ground after wakening, and recording the number and identities of all birds present. These roost counts were then used to estimated the number of birds at each site, as well as the percentage of marked birds. As individuals regularly visit other different sites, the number of birds marked at one specific site can be larger than the number of individuals roosting at this location. P: paintmarked; W: wingtagged; R: recognizable feature. For Northbridge, the number between brackets represent the number and percentage of marked individuals of the satellite roost.

| Roosting site (abbreviation) | GPS location | Number of marked birds | Roost size | Percentage marked |
|---|---|---|---|---|
| Balmoral Beach (BA) | -33.828494, 151.253983 | 209P, 12W, 6R | 88-165 | 94-96% |
| Clifton Garden (CG) | -33.841444, 151.252889 | 86P, 12W, 1R | 42-111 | 92-95% |
| Northbridge (NB) | -33.817167, 151.221694 | 78P, 2W, 4R | 56-72 (+21-27) | 93-98 % (100%) |

Table S2: Aggressive behaviours by SC-cockatoos recorded in the study, and their definitions

| Category | Behaviour | Definition |
|---|---|---|
| | Slow approach | Aggressor walks towards the receiver, with its head held high (Hardy, 1965; Levinson, 1980) |
| | Rushing | Aggressor runs towards its opponent; its head and neck are extended (Hardy, 1965; Power, 1966) |
| | Flight approach | Aggressor flies directly at the other individual (Hardy, 1965; Power, 1966) |
| Non-contact threats | Stare | Individual freezes, does not move while staring at another individual |
| | Beak-threat | Aggressor opens and closes its beak while making pecking a motion towards opponent, but does not make contact |
| | Beak-gape | Aggressor opens the beak while facing the opponent, but does not make contact (Hardy, 1965; Power, 1966; Garnetzke-Stollmann and Franck, 1991) |
| Displays | Crest-display | Crest is erected and displayed towards one (or several) other individual(s) |
| | Wingflapping | Wings are extended to either side of the body, while the aggressor faces the opponent (Power, 1966) |
| Contact threats | Bite | Aggressor's beak closes on some part of receiver's body (Hardy, 1965; Power, 1966; Garnetzke-Stollmann and Franck, 1991 |
| | Bill touch | Short bouts during which the beaks touch (Garnetzke-Stollmann and Franck, 1991) |
| | Foot on | Placing one foot on the recipients body, usually the tail |

( To be continued)

| Category | Behaviour | Definition |
|---|---|---|
| | Turn-towards | Individual suddenly turns towards the other individual preceding aggressive interaction. Often associated with beak-gape or beak-threat |
| | Fight | Both individuals fight. Can be aerial or on the ground. |
| | Displace | Individual displaces its opponent |
| | Crouch | Receiver of the aggression crouches (Seibert and Crowell-Davis, 2001) |
| | Retreat | Accelerated movement away from the opponent ( $>3$ body lengths) |
| | Move aside | Slight movement away from the opponent ($<3$ body-lengths) |
| | Fly away | Individual flies away, ending the aggressive interaction. |

( To be continued)

Table S3: Uncertainty of SC-cockatoos social hierarchies measured at each of the three roost sites. Correlations above 0.8 (repeatability by randomisation) and 0.5 (repeatability by splitting) suggest robust hierarchies (Sánchez-Tójar et al., 2018)

| Roosting site | Repeatability by randomisation | Repeatability by splitting (CI) | N individuals | N interactions |
|---|---|---|---|---|
| BA | 0.96 | 0.80 (0.75-0.85) | 126 | 1,737 |
| CG | 0.97 | 0.88 (0.84-0.91) | 93 | 2,735 |
| NB | 0.97 | 0.77 (0.70-0.84) | 74 | 1,005 |

Table S4: Density of aggressive networks, recorded either at group level, or subsetted to within sex interactions

| Roosting site | Group level | Males | females |
|---|---|---|---|
| BA | 0.22 | 0.52 | NA |
| CG | 0.64 | 0.80 | 0.65 |
| NB | 0.19 | 0.39 | NA |

Table S5: Correlations between SC-cockatoo hierarchies recorded at each roosting site during the two observation periods (July & September 2019)

| Roosting site | Correlation (CI) | N individuals | N interactions (July/September) |
|---|---|---|---|
| BA | 0.70 (0.41-0.86) | 24 | 88/189 |
| CG | 0.90 (0.84-0.95) | 40 | 672/805 |
| NB | 0.88 (0.53-0.98) | 9 | 16/55 |

Table S6: Number of SC-cockatoo cyclic triads compared to the total number of complete triads.

The number of each type of triads was determined on an outcome matrix, using igraph (Csardi and Nepusz, 2006).

| Roosting site | Overall | Males | Females |
|---|---|---|---|
| BA | 107/2168 (3%) | 26/393 (6%) | 0/8 (0%) |
| CG | 176/6850 (3%) | 39/1350 (3%) | 6/91 (6%) |
| NB | 34/1062 (3%) | 2/170 (1%) | / |

Table S7: Transitivity of aggression networks at SC-cockatoo group level, calculated for all individuals present at one site (overall), or within each sex.

| Roosting site | Overall | Males | Females |
|---|---|---|---|
| BA | 0.80 | 0.74 | / |
| CG | 0.90 | 0.89 | 0.74 |
| NB | 0.88 | 0.96 | / |

(a) (b)

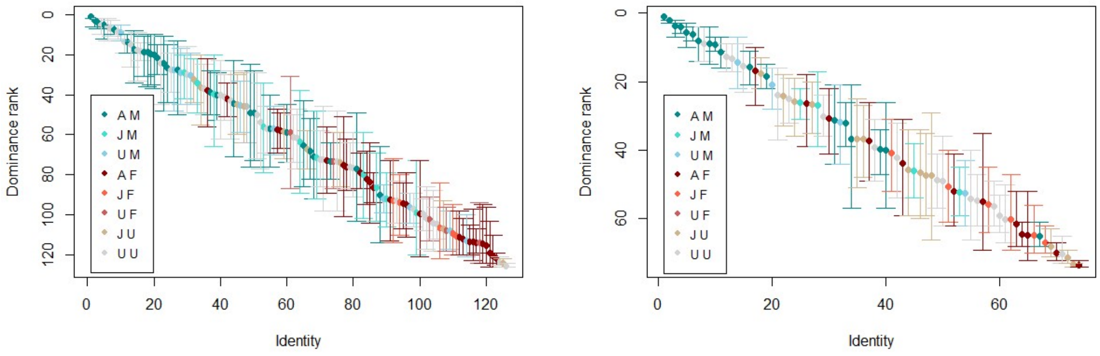


Figure S1: Hierarchies of two groups of sulphur-crested cockatoos, coloured by roost membership (left column), and their associated steepness (right column). a) Hierarchy of the BA roosting community, b) steepness of the hierarchy in the BA roosting community, c) hierarchy of the NB roosting community, d) steepness of the hierarchy in the NB roosting community. A: adult, J: juvenile, F: female, M: male, U: unknown (can be either unknown age or unknown sex)

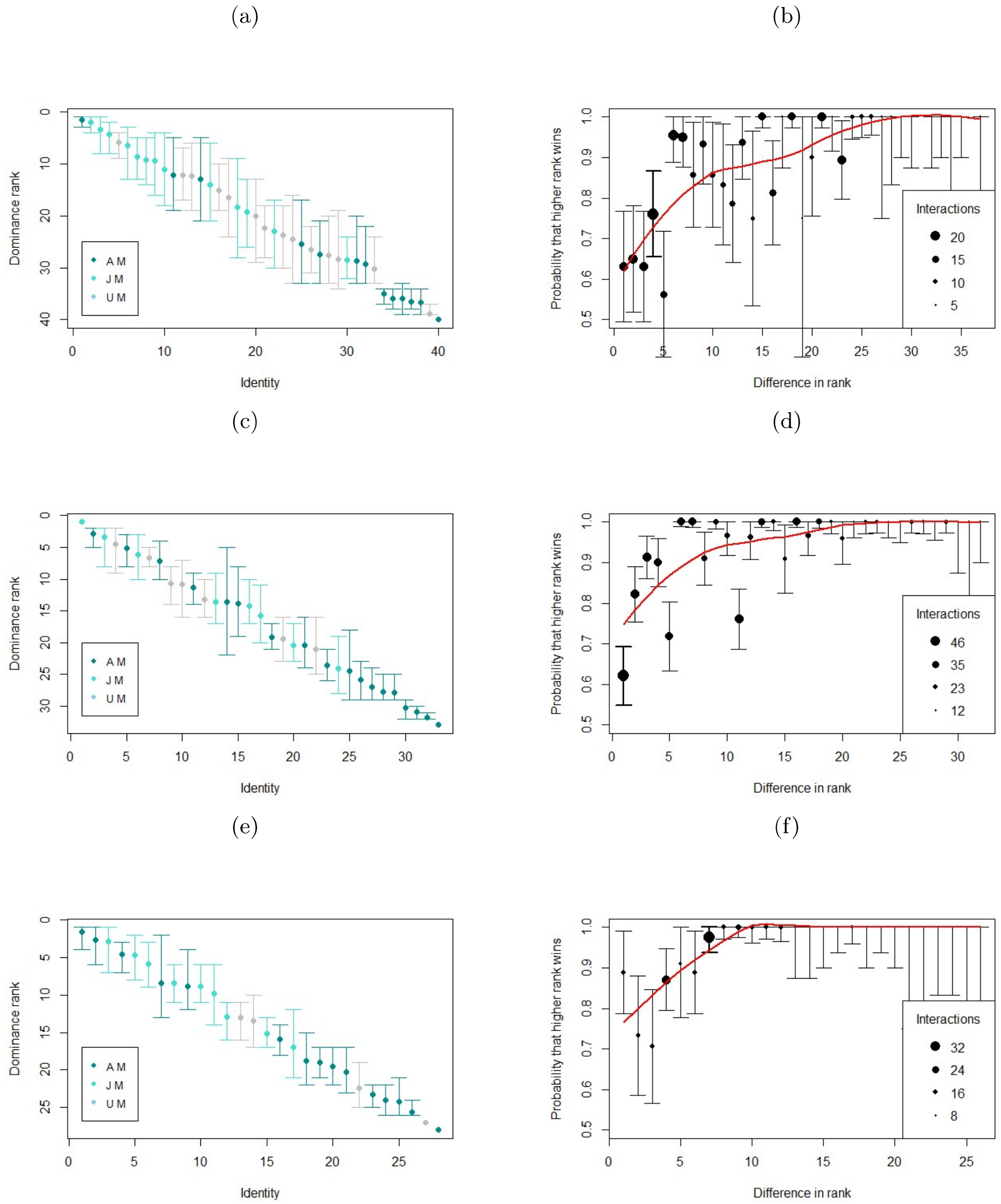


Figure S2: Hierarchies (left column) and their respective shapes (right column) of the hierarchies of three groups of sulphur-crested cockatoos, including only males. The dots are coloured by the age class of the individual. A M: adult male, J M: juvenile male, U M: male of unknown age: a) hierarchy of the male hierarchy at the BA roosting community, and its associated shape (b). c) Hierarchy of the male hierarchy at the CG roosting community, and its associated shape (d). e) Hierarchy of the male hierarchy at the NB roosting community, and its associated shape (f).

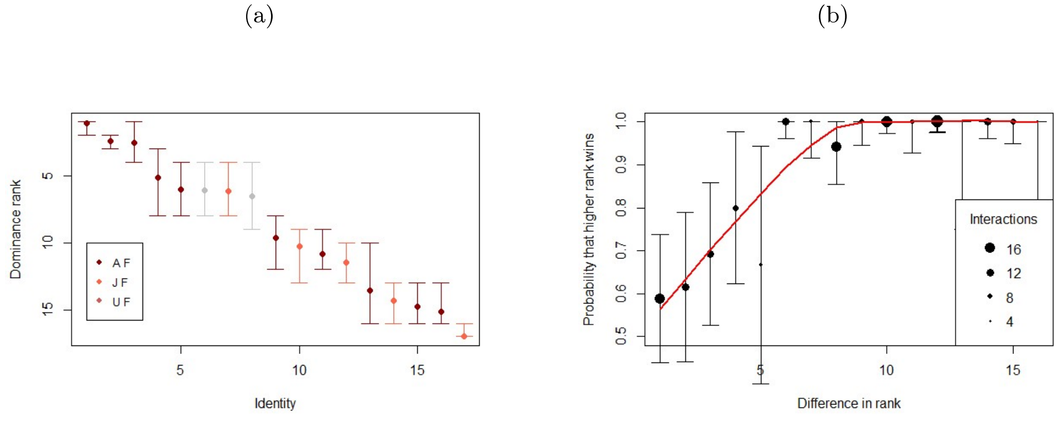


Figure S3: Hierarchy of female sulphur-crested cockatoos, at the CG roost site (a), and its associated shape (b).
The dots are coloured by age. A F: adult female, J F: juvenile female, and U F: female of unknown age.

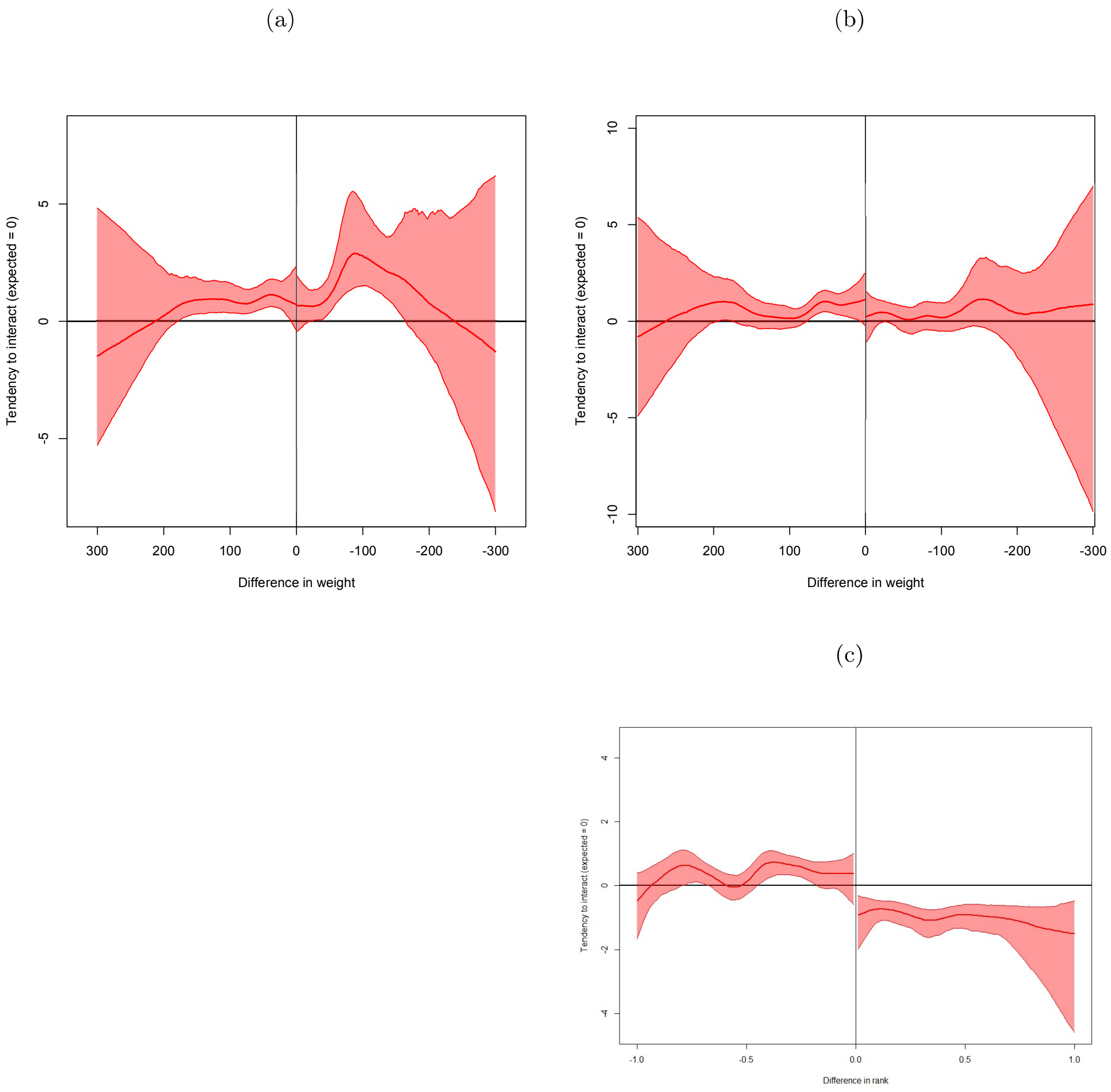


Figure S4: Tendency to initiate aggressive interactions, plotted by weight difference (top row - a, b) or rank difference (bottom row - c), depending on whether the initiator was considered not knowledgeable (left column) or knowledgeable (right column) of the social environment at the site of interaction. The thick line on each graph represents the median tendency to interact. The shaded areas represent the 95% confidence interval of the estimated interaction tendencies. If the tendency to interact follows a random pattern, the confidence intervals overlap with zero. A positive or negative value indicates that interactions between individuals of this specific rank difference occurred more or less often than expected by chance.
